# araCNA: Somatic copy number profiling using long-range sequence models

**DOI:** 10.1101/2025.01.20.633608

**Authors:** Ellen Visscher, Christopher Yau

## Abstract

Somatic copy number alterations (CNAs) are hallmarks of cancer. Current algorithms that call CNAs from whole genome sequenced (WGS) data have not exploited deep learning methods owing to computational scaling limitations. Here, we present a novel deep-learning approach, araCNA, trained only on simulated data that can accurately predict CNAs in real WGS cancer genomes. araCNA uses novel transformer alternatives (e.g Mamba) to handle genomic-scale sequence lengths (∼1M) and learn long-range interactions. Results are extremely accurate on simulated data, and this zero-shot approach is on par with existing methods when applied to 50 WGS samples from the cancer genome atlas. Notably, our approach requires only a tumour sample and not a matched normal sample, has fewer markers of overfitting, and performs inference in only a few minutes. araCNA demonstrates how domain knowledge can be used to simulate training sets that harness the power of modern machine learning in biological applications.

## Introduction

Somatic copy number alterations (CNAs) are genomic regions that are amplified or deleted when somatic cells replicate. They are a hallmark of many cancers and a driving factor of tumorigenesis (Zhang and Pellman, 2022), driving the amplification of oncogenes (Kim et al., 2020; Rosswog et al., 2021). CNA abundance is known to be associated with disease stage, prognosis and response to treatment, and particular cancer types can be characterised by specific classes of structural variation, often in specific chromosomal regions (Hadi et al., 2020). Accurate CNA profiles of cancer samples are important for downstream analysis such as association studies to identify underlying cancer signatures or as a prognostic biomarker (Steele et al., 2022).

The copy number landscape of a tumour can be profiled using array- and sequencing-based technologies where increases and decreases in signal intensity or sequence read depth correspond to gains and loss of genomic segments (Nakagawa et al., 2015; Pinkel and Albertson, 2005). Allelic information from single nucleotide polymorphisms (SNPs) can also be exploited to determine copy number (see **Figure 1, Figure 2**). Data can then be processed using CNA calling algorithms to derive copy number states.

**Figure 1:**
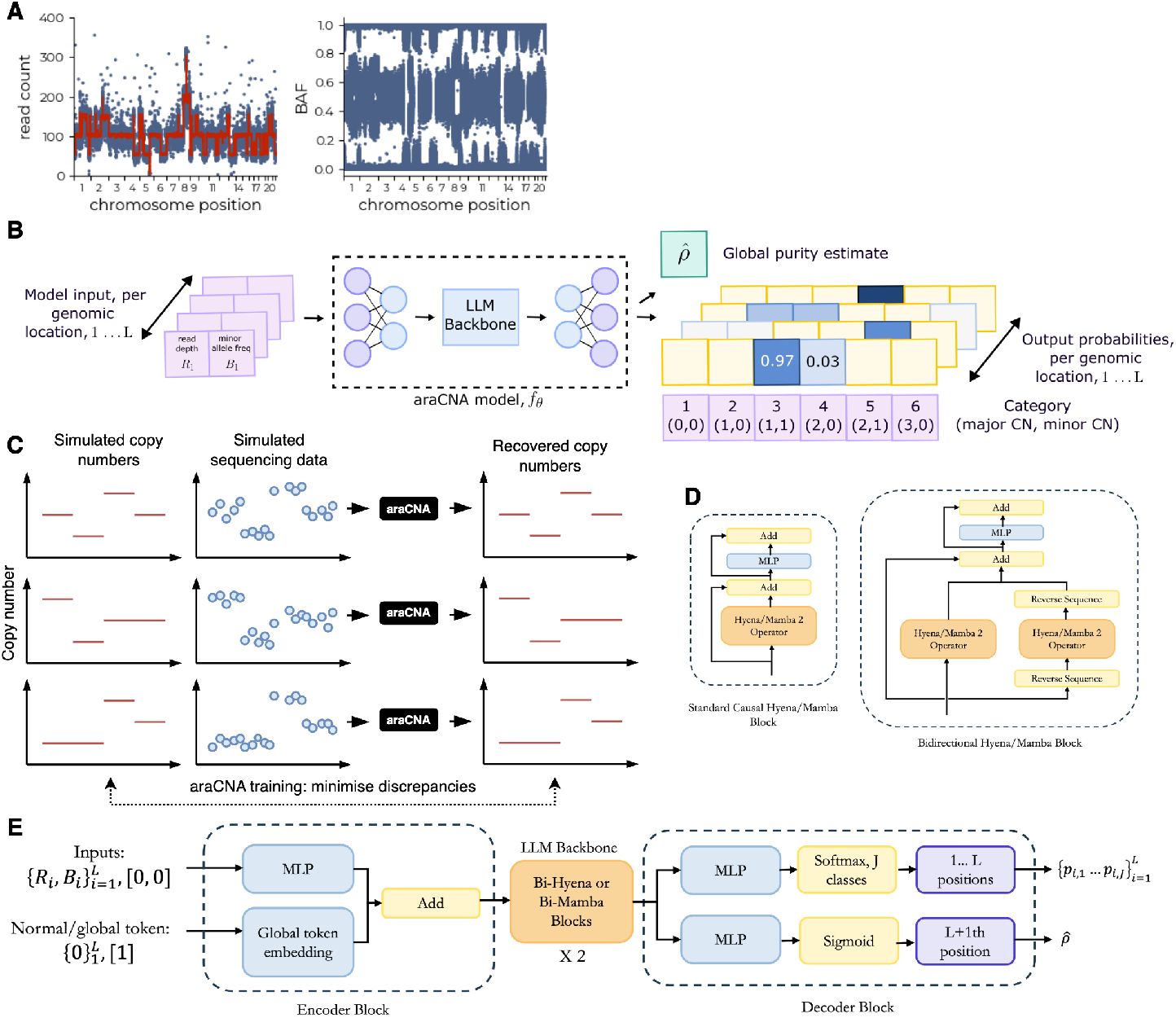
Overview of araCNA model. The (A) input data (read count and B allele frequency) is (B) converted by `araCNA` into a sequence of probabilistic copy number calls and global parameter estimates. (C) `araCNA` is trained using simulated data. (D) Bi-directional variants of the causal Hyena or Mamba blocks are used in the (E) `araCNA` model architecture.

**Figure 2:**
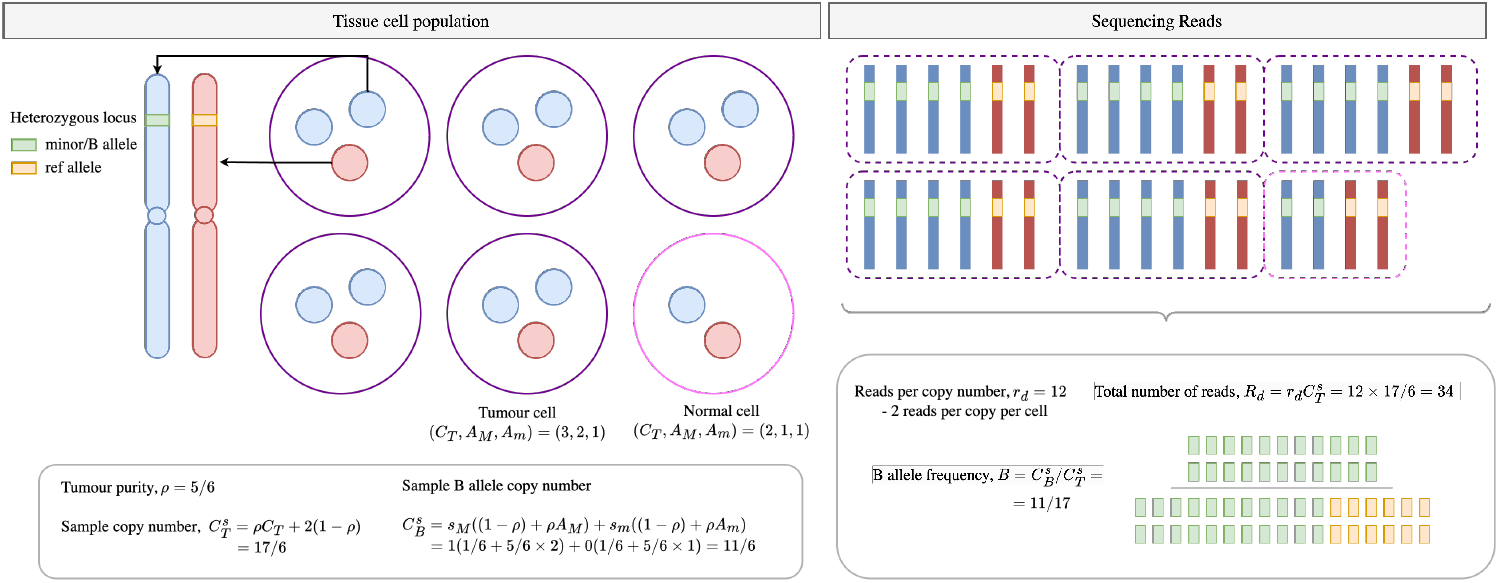
Mathematical construction of copy number calling. Illustration of how purity, copy number, read depth per-copy number and heterozygous loci result in measured read depth and B allele frequency.

CNA calling algorithms generally consist of an approach to segment the genome into regions of constant copy number and a copy number calling or classification step (into deletion, duplication, etc). Methods adopt a variety of approaches. This includes calling total copy numbers only (Chiang et al., 2009; Ivakhno et al., 2010; Ha et al., 2012; Xi et al., 2016) while others provide allele-specific major and minor copy number calling capabilities (Van Loo et al., 2010; Nik-Zainal et al., 2012). Others adopt a hybrid approach, first calling total copy numbers and then assigning these to major/minor copy numbers (Boeva et al., 2012; Talevich et al., 2016). Approaches that account for tumour heterogeneity can also provide sub-clonal copy numbers (Carter et al., 2012; Oesper et al., 2013; Fischer et al., 2014; Ha et al., 2014), again sometimes calling allele-specific copy numbers (Shen and Seshan, 2016; McPherson et al., 2017; Cun et al., 2018; Zaccaria and Raphael, 2020). When multiple tumour samples are available from the same individual, information from across samples can be used to posit evolutionary trees upon which somatic point mutations and CNAs can be placed (Satas et al., 2021; Myers et al., 2024).

Standard CNA callers adopt the use of segmentation algorithms or hidden Markov models which are effective at processing one cancer sequencing sample at a time but the low information capacity of these methods or models mean they cannot *learn* (Janiesch et al., 2021; Mohamed et al.). CNA callers process every sample as though it were the very first sample they have seen. Therefore, despite large-scale mapping exercises such as The Pan-Cancer Genome Atlas studies (Steele et al., 2022), few new CNA calling approaches have been developed.

While many areas of ‘omics analysis have been heavily influenced by deep learning in recent years, the technology has had relatively little impact on CNA calling algorithms. For example, the use of transformer-based models has led to the creation of massive foundation models for single-cell and integrative ‘omics (Szalata et al., 2024). However, the quadratic scaling limits of transformers have meant that these same principles are often not applicable to genome-based applications, like CNA calling, where distances between genomic regions of interest might be vast (Consens et al., 2023). One of the few examples is ECOLE, (Mandiracioglu et al., 2024), which is designed for whole exome sequencing (WES), which uses a transformer architecture to classify exome regions into neutral, deletion, and duplication, etc. Since WES has lower data resolution as it captures only the coding portion of the genome, ECOLE tackles the simpler task of classifying exome regions into these broad categories and does call the exact copy number, or allele-specific copy numbers.

In this paper, we propose a novel deep learning-based approach to the CNA calling problem, which we call `araCNA`, and is based on the use of recently developed transformer alternatives that provide genome-scale modeling capabilities. While existing copy number callers can be considered to be complex signal segmentation algorithms, `araCNA` *learns* to call copy number alterations and other features of sequencing data obtained from cancer samples. We describe how to train such a model and provide empirical evidence of its performance. The approach we suggest therefore opens up new opportunities for creating CNA callers that learn and can be fine-tuned for specific sub-tasks of interest, or integrated directly with deep learning models for other data modalities.

## Materials & Methods

### Model overview

The input data for `araCNA` is assumed to be a sequence of allele-specific read counts at genomewide loci (see **Figure 1A**) which can be converted into total read depth (*R*) and B allele frequency (*B*) values as is commonplace for copy number callers. These are fed into a longrange sequence model which converts the input sequence into an output sequence consisting of the probabilities of major and minor copy number values at corresponding loci. Global parameters of interest such as tumour purity or ploidy can also be provided (**Figure 1B**).

### Mathematical Preliminaries

We first define the mathematical construction, similar to that first shown in Van Loo et al. (2010). We define *C*_*T*,*i*_ as the total copy number at locus *i* in the tumour, *C*_*P*,*i*_ as the copy number of the paternal chromosome at locus *i, C*_*M*,*i*_ as the copy number of the maternal chromosome at locus *i* and *C*_*B*,*i*_ is the B allele copy number. We further define *ρ* as the purity of the tumour sample (the proportion of tumour vs non-tumour) and *r*_*d*_ as the expected number of sequencing reads per copy number. In practice, samples are sequenced up to a given average read depth, which assumes a uniform coverage of the short reads across the whole genome, (Sims et al., 2014). When many duplicated regions exist, the actual average read depth per copy number *r*_*d*_ is unknown. Finally, we assign *R*_*i*_ to mean the total number of reads at a locus and *B*_*i*_ as the B allele frequency (BAF). The sequence data is therefore a collection 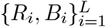 for *L* loci.

For a pure tumour sample, we have the total copy number as

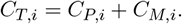

Considering sample impurity, we define the sample copy number 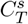 as

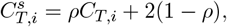

where we assume the contaminating normal cells have copy number 2 at all loci. While normal cells may possess some copy number variants, the size of these regions are typically negligible compared to the cancer-associated alterations we aim to detect and so we ignore these for simplicity.

The B allele copy number is defined as:

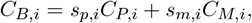

where (*s*_*p*,*i*_, *s*_*m*,*i*_) ∈ *{*(0, 0), (0, 1), (1, 0), (1, 1)*}* denotes whether the paternal and maternal chromosomes respectively have the specified SNP B allele at locus *i*.

Adding sample impurity we have:

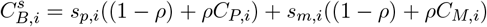

The total number of reads at a locus *R*_*i*_ are then:

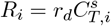

While the B allele frequency (BAF) *B*_*i*_ at locus *i* is given by:

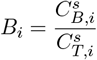

Both *R*_*i*_ and *B*_*i*_ are measured data, obtained from sequencing a tumour sample. Figure 2 illustrates the relationship from (*ρ, r*_*d*_, *{C*_*P*,*i*_, *C*_*M*,*i*_*}*) → *{R*_*i*_, *B*_*i*_*}*. Our aim is to do the reverse-to infer (*ρ, r*_*d*_, *{C*_*P*,*i*_, *C*_*M*,*i*_*}*) from *{R*_*i*_, *B*_*i*_*}*. However, we can see in the above formulation, that both *R*_*i*_ and *B*_*i*_ remain the same if we were to swap the values for *C*_*P*,*i*_ and *C*_*M*,*i*_, meaning that we are unable to infer the parental copy numbers they are non-identifiable. We can instead introduce, *A*_*M*,*i*_, *A*_*m*,*i*_ = max(*C*_*P*,*i*_, *C*_*M*,*i*_), min(*C*_*P*,*i*_, *C*_*M*,*i*_) as the major/minor allele-specific copy numbers, which are identifiable. Hence, we amend our aim to infer (*ρ, r*_*d*_, *{A*_*M*,*i*_, *A*_*m*,*i*_*}*) from *{R*_*i*_, *B*_*i*_*}*.

### The araCNA model

The function of `araCNA` can be summarised as 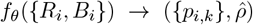 where *f*_*θ*_ is the long-range sequence model parameterised by network weights *θ*. 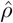 is `araCNA`’s global purity estimate. While *p*_*k*,*i*_ is the probability that `araCNA` assigns the locus as belonging to copy number profile *K*_*j*_. The profile categories *K*_1_, …, *K*_*J*_ correspond to major/minor parental copy number combinations, with *K*_1_ := (*A*_*M*_ = 0, *A*_*m*_ = 0), *K*_2_ := (*A*_*M*_ = 1, *A*_*m*_ = 0) etc. Here, 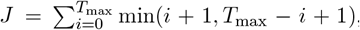, where *T*_max_ is a hyper-parameter corresponding_max max_ to the maximum modelled total parental copy number. From these, we can also estimate 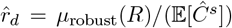, where *µ*_robust_(*R*) is the robust or trimmed mean of the read depth vector and 𝔼 [*Ĉ*^*s*^] is the expected value of the overall sample copy number (ploidy).

We trained our model using simulated datasets where the ground truth copy numbers and purity are known. The details of this simulation are discussed later in this section. The loss function consists of a supervised sequence loss and a supervised global loss. The supervised sequence loss is the cross entropy:

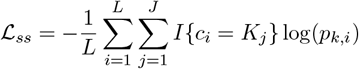

where *L* is the sequence length, *c*_*i*_ ∈ *K*_1_ … *K*_*J*_ is the known target profile of a genomic locus. The supervised global parameter losses are:

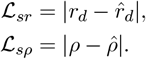

The total loss is then given by:

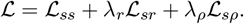

We found *λ*_*r*_ = *λ*_*ρ*_ = 1 to work well. The model was trained by iteratively increasing the complexity of the simulated data.

### Model Architecture

Sequence inputs are entered into `araCNA` together with global placeholders, and projects these through an encoder block, backbone and decoder block to give the copy number and purity outputs, **Figure 1C**. Elements used as global estimates (i.e. purity) in the output are appended to the end of the normal sequence, with zero values for read depth and minor allele frequency.

To differentiate these elements from normal elements, we included a global token feature, encoded as 0 for normal elements and 1 for global prediction sequence elements. The normal observational data (read depth and minor allele frequency), is projected into a dimension *d* using a multi-layer perception (MLP) with ReLU non-linearities, while the token feature is projected into dimension *d* using a simple embedding. The two inputs are combined in this projected space, to serve as the encoder block for the main backbone of the model. The output from the main backbone diverges into two streams in the decoder- one for prediction of the copy numbers, and one for prediction of the global parameters. Both streams use MLPs with ReLU activation, with a softmax to project the copy number stream into a probability over the *J* possible classes, and a sigmoid to project the purity estimate between 0 and 1.

We created two implementations of `araCNA` which use two recently developed long-range sequence models, Hyena (Poli et al., 2023) and Mamba (Gu and Dao, 2024; Dao and Gu, 2024), as part of its architecture. These long-range sequence models known as state space like models (SSMs), were developed to achieve similar results to transformers but avoid using the computationally expensive self-attention mechanism, where the number of operations scales quadratically with sequence length and therefore is prohibitive for genome-scale data (Consens et al., 2023). In contrast, Mamba and Hyena-based models offer training times that scale linearly with sequence length. We call the `araCNA` models with Mamba/Hyena blocks `araCNA`-mamba and `araCNA`-hyena respectively. Mamba-2 is the more popular of the SSM models, but being constrained to GPUs with Nvidia A100 architecture (for training and inference). We adapt the blocks to be bidirectional rather than causal (unidirectional) as used in natural language processing (**Figure 1E**).

The hyperparameters of `araCNA`-mamba and `araCNA`-hyena are outlined in the **Supplementary Information**, resulting in parameter counts of 70K and 12.09M respectively. Each hyena block contains two positional encodings of length *L*_max_ corresponding to a hyperparameter of the maximum input sequence length. For our models, *L*_max_ was set to 1M, hence the majority of the parameters, 12M, can be attributed to the positional encodings. We note that 70K parameters for `araCNA`-mamba is relatively small for a modern neural network, especially given the task at hand is not trivial (Habib and Qureshi, 2022).

### Smoothing

Given the noise in real data, many copy number calling methods employ smoothing or thresholding in some way to reduce the number of CNA segments and reduce false positives. For example, ASCAT has a ‘penalty’ parameter that controls smoothing (Van Loo et al., 2010), Battenberg has several ‘gamma’ hyperparameters (Nik-Zainal et al., 2012), HMMCopy has ‘e’ and ‘strength’ parameters (Ha et al., 2012) and CNV Kit uses thresholding to segment and call different copy numbers (Talevich et al., 2016). We also employ a smoothing technique, based on the predicted output probabilities.

Here, we wish to find the ‘best’ joint sequence **S** ∈ *{*1, … *J }*^*L*^ given the model output probabilities, *p*_*k*,*i*_, that a locus, *i*, is in copy number state *k*. We can find the optimal sequence by balancing the probabilistic evidence of being in a given state, with the penalty *λ*_*t*_ of transitioning to a different state as:

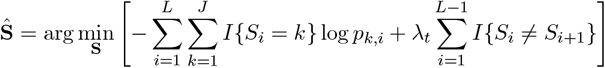

For a given *λ*_*t*_, we solve this using a dynamic programming approach similar to the Viterbi algorithm used in HMMs for finding the most likely sequence of hidden states. We found *λ*_*t*_ = 500 to be suitable.

### Synthetic data simulation

We generated synthetic copy number profiles using the following procedures:

1. *Sampling the number of segments*. We sample the approximate number of copy number segments, 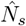 using a mixture approach; first, we sample a uniform variable, *u* such that under a user-defined swap probability, *q*_*s*_, the number of segments is sampled uniformly between 1 and *N*. When *u > q*_*s*_, a Poisson distribution is used to skew sampling towards smaller total segments. This is to oversample harder cases with fewer segments where it is harder to estimate global parameters like read depth per copy number and purity.
2. *Sampling the segment breakpoints*. This is done by randomly sampling 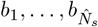 breakpoints from 1 … *L*, the unique set of these breakpoints defines the segments, and 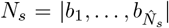. We only keep the segments that have a minimum segment length of *L*_min_.
3. *Sampling the segment profiles*. We sample *A*_*M*_, *A*_*m*_ of each segment from the possible copy number profiles. We inject logic here to preferentially sample profiles closer in copy number to 1-1 when there are fewer segments. This is due to the identifiability issue. When there are more segments, profiles are sampled more uniformly but still with a preference for lower copy numbers, to inject an implicit bias towards lower ploidy solutions when the model is unsure.

From a sampled profile, we simulate the sequencing read depth and B allele frequency data. Each of the *L* loci is considered a commonly varying single nucleotide polymorphism, SNP. For both parental alleles, *A*_*M*_, *A*_*m*_, we sample each SNP as binomial with a probability of 0.5, we also sample the purity, *ρ*, uniformly from a range between 0.5 and 1. This gives the sample minor allele copy number, 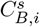 and the sample total copy number, 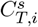. The read depth per copy number, *r*_*d*_ is sampled uniformly between 5 and 70, and together with 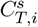 the overall read depth, *R*_*i*_ is sampled from this mean with additional noise. The BAF, *B*_*i*_ is sampled using total reads sampled based on *R*_*i*_ and the subset of B-allele reads using a binomial probability of 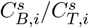, with added noise.

In real data, there exist regions of prolonged homozygosity that can be attributed to identityby-descent (IBD) regions (i.e identical regions inherited from both parents due to a common ancestor), Figure 3. To emulate this, we also randomly inject regions of prolonged homozygosity into the model. In these IBD regions, the BAF cannot be used to infer copy-number, and the model must use context from before/after the homozygous region for correct prediction.

**Figure 3:**
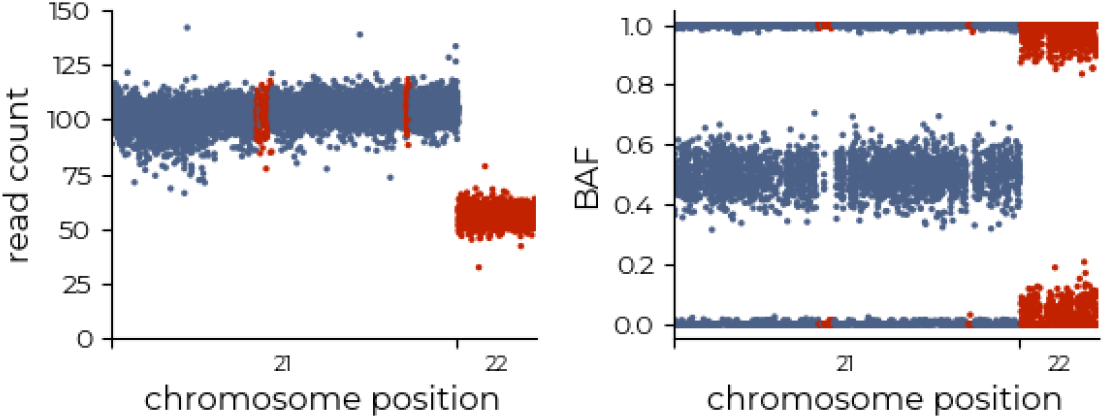
Real sample example illustrating prolonged homozygous regions on chromosome 21, where the BAF noise is very low. In contrast, the region on chromosome 22 indicates a minor copy number of 0, and hence although the BAF profile is close to 1 and 0, the noise profile is larger, where normal contamination at heterozygous loci brings the BAF closer to 0.5. Red colouring indicates regions of interest.

From this sampling procedure, we therefore have a set of targets (*{A*_*M*,*i*_, *A*_*m*,*i*_*}, ρ, r*_*d*_) that generate inputs *{R*_*i*_, *B*_*i*_*}*, which together are used in the training of `araCNA`. Hence, `araCNA` can be interpreted as performing inference on the above statistical approach, when (*{A*_*M*,*i*_, *A*_*m*,*i*_*}, ρ, r*_*d*_) are treated as unknowns. Further details of this procedure are included in **Supplementary Information**.

### Training Procedure

For this work, we trained `araCNA` using simulated copy number profiles. We elected to follow this approach since (i) there exists no ground-truth, high-resolution copy number profiles upon which `araCNA` could be trained and (ii) we avoid using copy number profiles produced by other methods in order to perform fair comparisons later on. For the simulations, we set the maximum considered total copy number *T*_max_ as a hyperparameter, we then do the following:

1. Sample the approximate number of segments, 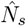 in the total sequence.
2. Sample the segment breakpoints, that gives the actual number of segments *N*_*s*_ in the total sequence.
3. Sample the copy number profiles for each segment, *A*_*M*,*i*_, *A*_*m*,*i*_ (model targets).
4. Sample the sequence data, *R*_*i*_, *B*_*i*_ (model input) across all segments, based on sampled purity, *ρ* read depth per copy number, *r*_*d*_ and copy numbers, *A*_*M*,*i*_, *A*_*m*,*i*_.

Noise and tumour purity parameters are sampled randomly during training allowing `araCNA` to see different sequence data distributions and learn how to process these into copy number calls. As a consequence, when used at test-time for zero-shot inference, it is not necessary to retrain `araCNA` to process any tumour sample unlike conventional CNA callers which need to be adapted to each sample every time.

To train `araCNA` we adopted an iterative warmup procedure, gradually increasing the complexity of the problem. We found this was necessary for the model to learn, and a similar approach was taken in Nguyen et al. (2023) with gradually increasing the sequence length.

The training procedure was:

1. Begin the synthetic data generating procedure with *ρ* = 1, and without sampling the noise parameters. Use only up to a maximum total copy number of 2, that is profiles, (*A*_*M*_, *A*_*m*_) ∈ *{*(0, 0), (1, 0), (1, 1), (2, 0)*}*. Sample *r*_*d*_, and start with sequence length 10000. Train until convergence.
2. Using the previously trained model weights as initialisation, add in purity and noise parameter sampling. Train until convergence.
3. Using the previously trained model weights as initialisation, slowly increase the maximum total copy number to 8. Train until convergence.
4. Using the previously trained model weights as initialisation, slowly increase the maximum sequence length to 650,000. Train until convergence.

### Metrics

We define the metrics used for reconstruction accuracy as follows. The BAF root mean square error (RMSE) is calculated as:

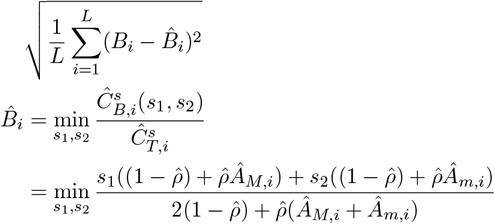

where the reconstructed BAF, 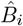 is calculated from the most probable haplotype (*s*_1_, *s*_2_) ∈ *{*(0, 0), (0, 1), (1, 0), (1, 1)*}* at each loci.

The read depth mean absolute error (MAE) is calculated as:

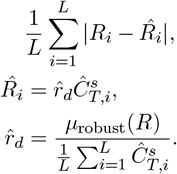

We opted to use absolute error here, due to mapping errors that can lead to read depth and hence RMSE inflation. Here, *µ*_robust_(*R*) is the robust or trimmed mean of the data, excluding data at the highest/lowest 5% of values.

Battenberg may assign a fraction of each segment to a subclone with a different set of copy numbers (Nik-Zainal et al., 2012). Hence, for the Battenberg multiclonal reconstruction, we account for the fraction attributed to some sub-clone, for that segment. This gives an extra set of parameters for each loci/segment; *τ*_*i*_, the fraction of the sample at that segment attributed to clone 1, 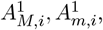, as well as (1 − *τ*_*i*_), 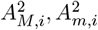 for clone B. We therefore adapt the reconstruction to follow the Battenberg formulation (Nik-Zainal et al., 2012), and in their implemented code (https://github.com/Wedge-lab/battenberg/), where the sample copy numbers for each locus is given by:

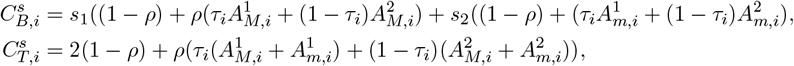

The analysis for the non-multiclonal Battenberg reconstruction uses the copy number predictions of the clone estimated as having the highest proportion (i.e max(*τ*_*i*_, 1 − *τ*_*i*_) at each locus, with the normal formulation.

The concordance, *CC* between two methods, 1 and 2, is calculated according to:

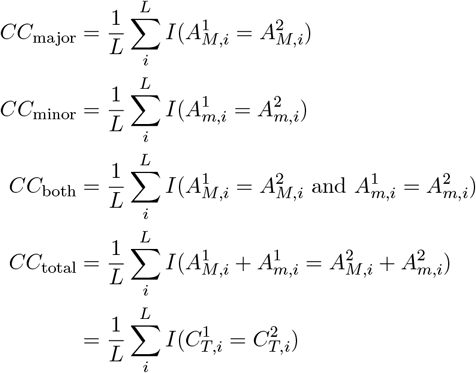

For methods like HMMCopy, only *C*_*T*_ is output, so only *CC*_total_ can be measured. When method 1 is the ground truth, like in simulations, this can be interpreted as an accuracy.

## Results

### Simulation Study

We first compared results from our two `araCNA` variants (`araCNA`-mamba and `araCNA`-hyena) using simulated data, sampled from the same generating procedure from which the training data was also sampled. We show results from 100 simulated test genomes, with a maximum sampled sequence length of 650k, to emulate the data size of a real SNP dataset. **Figure 4A** shows both models achieve high copy number classification accuracy for the task though `araCNA`-mamba slightly outperforms `araCNA`-hyena. Since in this case, we know the ground truth, we can directly measure accuracy. However, in real data, there is no known ground truth, so we also include the reconstruction error for the two models. **Figure 4C** shows unsupervised metrics for performance, where better methods are likely to have a lower BAF and read depth reconstruction. When examining tumour ploidy and purity estimates (**Figure 4B**) and reconstruction performance (**Figure 4C**), the differences are small though `araCNA`-mamba does achieve slightly better metrics than `araCNA`-hyena. **Figure 4D,E** show the reconstructed read depth and BAF and predicted copy numbers from each model for one of the example simulated genomes.

**Figure 4:**
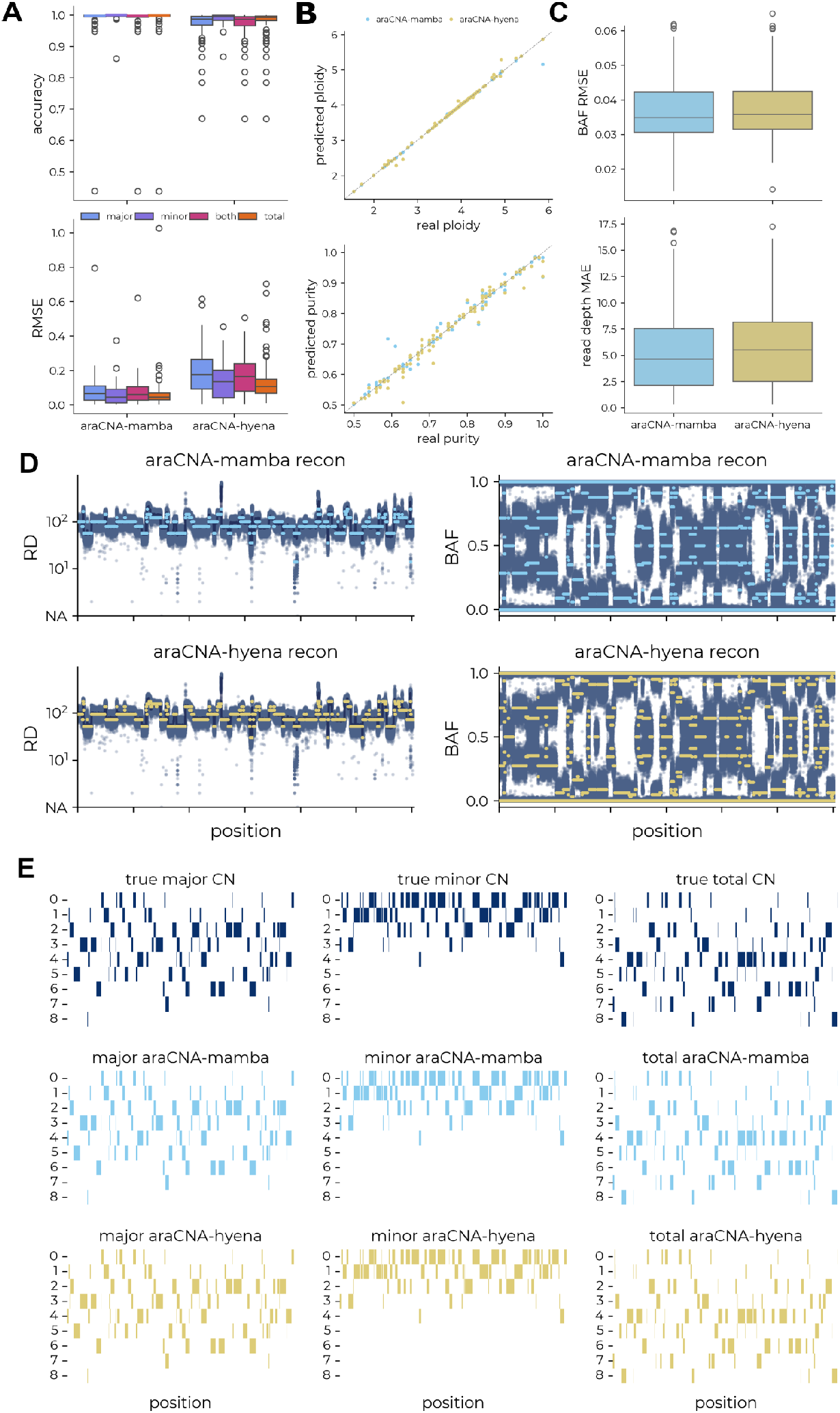
Results for araCNA model variants on simulated data. A-C show aggregated results across 100 simulated test genomes. (A) The distribution of concordance and RMSE between the predicted and true copy numbers. (B) The predicted ploidy and purity against the true purity and ploidy. (C) The distribution of mean reconstruction error: BAF root mean squared error (RMSE) and read-depth mean absolute error (MAE). (D) The reconstructed read depth and BAF and E) the underlying predicted copy number segments for `araCNA`mamba and `araCNA`-hyena on an example simulated test set.

The Hyena and Mamba models were trained using the same underlying simulation and training procedure. Differences in results can therefore be traced to differences in the model architecture or sensitivity to hyperparameters. We found that Hyena often took longer to converge during training, becoming stuck in local minima, likely due to the size of the model (12M parameters). In contrast, the Mamba model (70K parameters) achieved better performance despite its significantly smaller size and need for an A100 GPU.

### The Cancer Genome Atlas

Next, we compared `araCNA` to a number of existing CNA calling tools with whole genome sequencing data from a selection of 50 tumour samples chosen from the colorectal (CRC), breast (BRCA) and ovarian (OV) cancer cohorts of The Cancer Genome Atlas (TCGA). We chose these three cancer types due to the extensive and well-characterised aneuploidy present in tumours of these types. We utilised two of the most well-used algorithms that can call allele-specific copy numbers: ASCAT, (Van Loo et al., 2010), and an adaption of ASCAT called Battenberg, (Nik-Zainal et al., 2012), which can also model sub-clonal populations, by optionally assigning a fraction of each CNA segment to another subclone. We compared results to Battenberg without and with its default sub-clonality modeling.

We also compared to CNVkit, (Talevich et al., 2016), which works with WGS although being intended primarily for WES and gives allele-specific copy numbers after first calling total copy numbers. Finally, we report results for HMMCopy, which only calls total copy number (Ha et al., 2012). Both ASCAT and CNVkit performed well in a recent benchmark (Masood et al., 2024), while Battenberg and HMMCopy are popular methods included in other method benchmarks (Zaccaria and Raphael, 2020; Alkodsi et al., 2015). Details of their implementation can be found in the **Supplementary Information**.

Since there are no ground truth copy number profiles for these tumours, we used proxy measures such as reconstruction error as in the simulation study to provide an unsupervised metric for performance, where better methods will be expected to have low reconstruction error. However, it is also important to consider both the number of segments and the range of modeled copy numbers produced by each CNA calling approach. A low reconstruction error accompanied by a large number of highly variable copy number segments may suggest overfitting, while high copy number calls at high read depth regions will have a lower reconstruction error but could be less plausible. **Figure 5** illustrates the analysis of a TCGA ovarian cancer sample using all methods.

**Figure 5:**
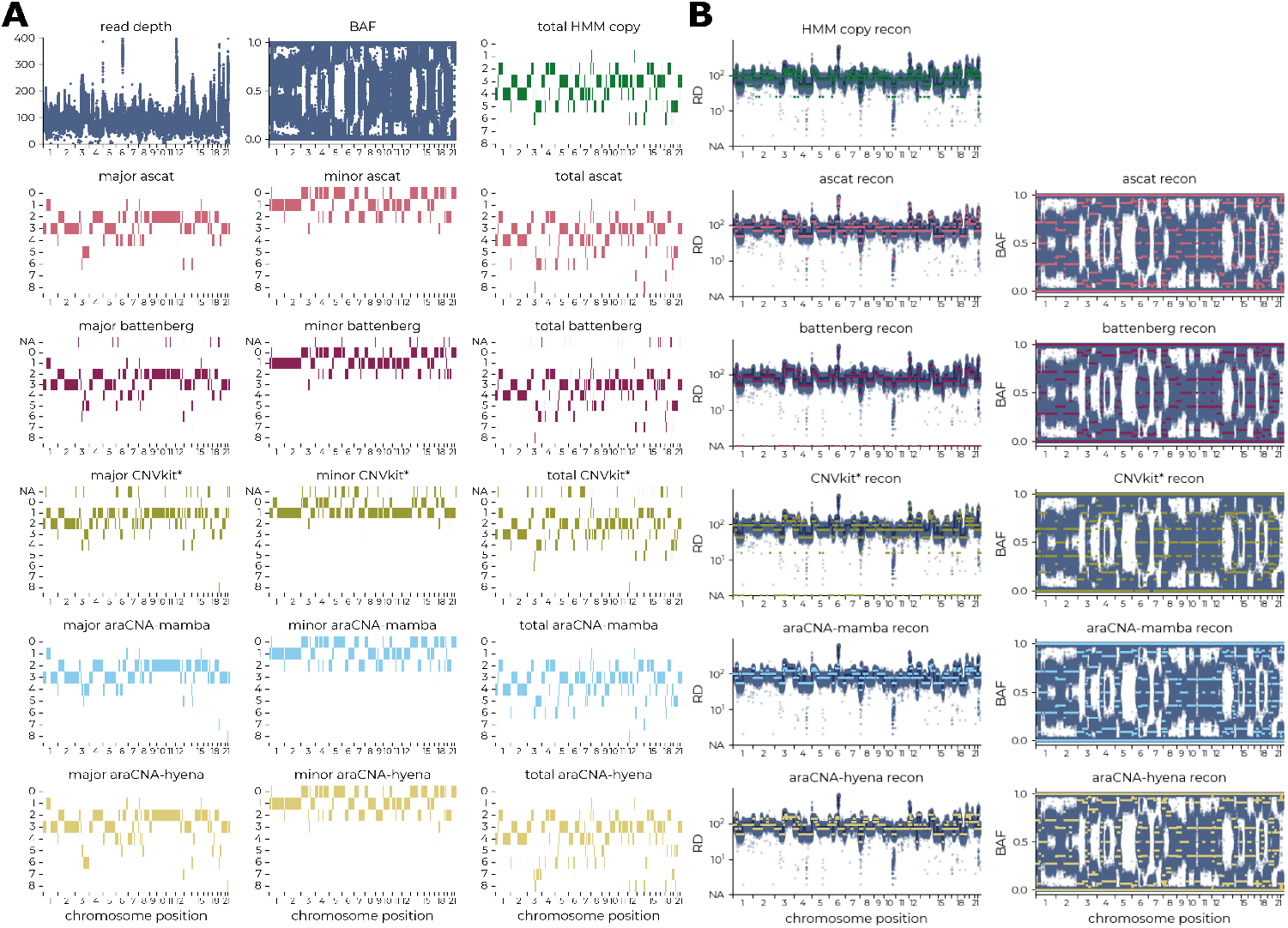
Representative TCGA ovarian cancer sample. Each row shows A) the called copy numbers and B) respective reconstruction of each caller, where `araCNA` has approx. 80% concordance with ASCAT, Battenberg and HMM Copy.

**Figure 6A** shows the distribution of the root mean-squared error (RMSE) and mean absolute deviation (MAE) for the reconstruction of the B allele frequency and read depth over the fifty samples. Using both `araCNA` variants for zero-shot inference of the copy number states gives comparable reconstruction performance to the existing CNA calling methods while using similar numbers of segments. While ASCAT and Battenberg are able to achieve slightly improved reconstruction error performance, when we examined the top five percent of total copy number calls produced, we found that ASCAT and Battenberg sometimes assigned as high as 100 copies to localised genomic regions. It is difficult to verify the correctness of such calls given the lack of ground-truth, however, we found most of these regions contained few or no known cancer or other functional genes (**Supplementary Figure SI-1**).

**Figure 6:**
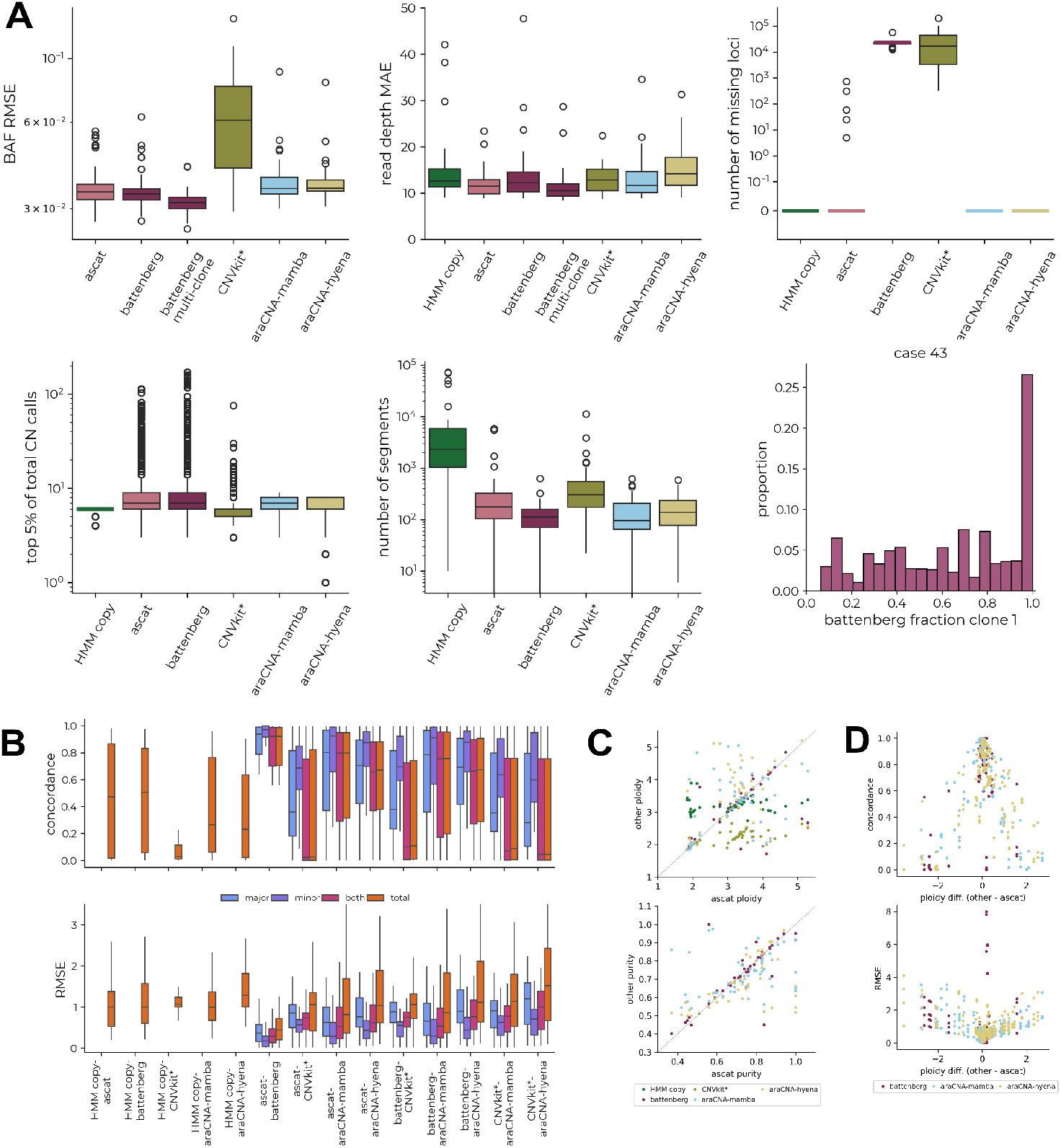
Comparison between CNA callers across 50 TCGA cancer samples. (A) Boxplots showing the B-allele-frequency root-mean-squared reconstruction error (RMSE), read-depth mean absolute error (MAE), number of missing loci after calling, copy number distribution of top 5% of copy number calls and the number of different copy number segments identified across the tumour samples. Example distribution of Battenberg clonal fraction for a particular tumour. (B) The distribution of concordance and RMSE between the copy number predictions of each method. (C) The predicted tumour purity and ploidy against the ASCAT-derived purity and ploidy. (D) Concordance and RMSE of predicted copy numbers against the ploidy difference between methods and ASCAT.

Further, Battenberg’s multi-clonal formulation uses the same input data with more degrees of freedom to model clonal fractions (see **Methods**). In the presence of clones, we would expect to see a concentration of fraction values around discrete modes, corresponding to the fraction of the sample originating from a subclone with a diverging copy number pattern across the genome.

We found that in most tumours there were no clear modes in the distribution of clonal fraction estimates, **Supplementary Figure SI-2**, indicating overfitting through this additional degree of freedom, to give the observed lower reconstruction error, **Figure 6A**.

We next looked at pairwise copy number classification concordance and the root mean squared difference between copy number calls from different methods (**Figure 6B**). `araCNA`-mamba achieves a median of 80% concordance with ASCAT for all copy number calls across the 50 TCGA cancer samples and a median of 70% concordance with Battenberg while `araCNA`-hyena was substantially less concordant with ASCAT and Battenberg. However, when the discrepancies are measured using root mean squared difference, the average differences were much less than one and we found that most discrepancies were due to small numerical differences (e.g. 1 ↔ 2, 2 ↔ 3, etc). When we further examined the differences between ASCAT and `araCNA`, we found that these appeared to be primarily driven by differences in major and total copy number calls in some samples.

To further investigate these discrepancies, we compared tumour ploidy and purity estimates using ASCAT as a baseline. While tumour purity estimates were well correlated between all methods, tumour ploidy estimates differed between methods for some tumours (**Figure 6C**). Interestingly, although `araCNA` is giving a different copy number profile for some samples compared to ASCAT, it still gives low reconstruction error. We surmised that for some tumours there is weak identifiability (see **Figure 7**) and `araCNA` identifies a different ploidy state to ASCAT but with a corresponding copy number profile that is still compatible with the observed data. Indeed when we examined the concordance between `araCNA` and Battenberg with ASCAT, we see a reduction when there is a significant departure from the ploidy state estimated by ASCAT (**Figure 6D**). Interestingly, where there are disagreements, tumours classified as near-triploid by ASCAT will generally be classified in a higher ploidy state by `araCNA` and vice-versa which is a hallmark of the identifiability issue. There also exists a subset of tumours where ASCAT and Battenberg disagreed on tumour ploidy despite their algorithmic similarities which further highlights sensitivity of methods to the identifiability issue.

**Figure 7:**
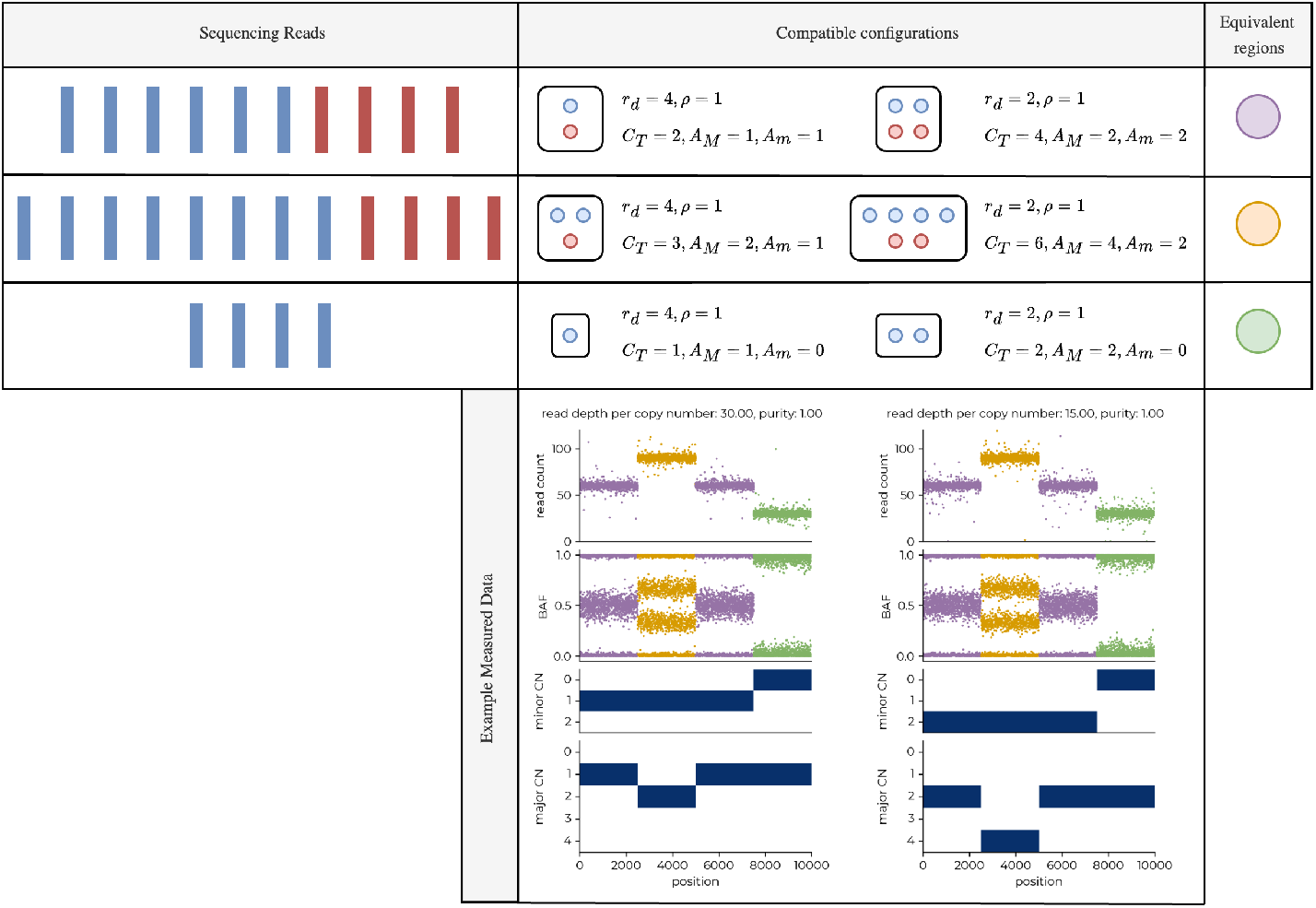
Identifiability of copy number profiles. Example illustrating how different copy number profiles can result in the same observed sequencing read depth and B-allele frequency measures. Here, each copy number profile on the right is exactly double that of the left, while the read depth per copy number value is half the left. Blue and red reads are those from the paternal/maternal chromosome respectively, where the B-allele frequency will reflect their chromosome ratio at heterozygous loci. This combination results in the exact same measured data, as highlighted in the bottom panel, where the colours link to each configuration profile above. Parameters are defined in **Methods**. In general, many combinations of copy number profile, tumour purity and read depth per copy number parameters may give rise to similar observed data. In the presence of noise, segments that may uniquely determine the copy number profile can be missed.

Overall, while `araCNA` was only trained with simulated data only and applied using zero-shot inference, it appears to perform as well as standard CNA callers on real tumour sequencing data with the caveat that there is a limit to our ability to quantify performance due to a universal lack of ground-truth data. This suggests that this form of

## Discussion

Our results have shown that it is possible to develop a novel deep-learning approach to predict copy number alterations from whole genome sequencing. We have built a training simulation framework to generate supervised training data for this biological problem that otherwise would have no known ground truth. We applied our model using zero-shot inference to TCGA cancer samples and found that `araCNA` achieves comparable performance to existing methods despite being trained only on simulated datasets. This highlights the utility of having strong mechanistic models that relate copy numbers to sequencing data. Overall, these results demonstrate how we can use domain knowledge to develop simulation training sets for simulation-based inference in biological applications, where computing the posterior may be intractable or computationally intensive (Cranmer et al., 2020).

We have taken advantage of recent advancements in deep learning, such as SSM-like models, that allow for extremely long sequence lengths, on the order of genomic data, and to learn longrange interactions for fast CNA inference. This approach is not limited to CNA calling, and could be repurposed for inference in many biological applications where simulations are already used. Our experiments with Mamba and Hyena highlight that the choice of architecture can be important and performance can vary between models for a given application.

Importantly, once constructed it is possible to further refine and retrain a model like with additional data. Unlike classic CNA callers, `araCNA` can continue to learn and improve using standard deep learning techniques such as fine-tuning or transfer learning. For instance, these approaches maybe used to adapt the model due to some distributional shift between the simulated training data used in training and the actual sequencing data. This might occur if, for example, there was additional sequencing noise and artefacts from sequencing of FFPE tumour samples.

While we limited ourselves in this work to training on simulated data, it is possible to use CNA profiles from existing copy number callers for training. However, since these would not be ground-truth data, this would result in a model which is an *emulator* of the existing copy number caller. While emulators could not normally exceed the original CNA caller in terms of performance, they can offer usability advantages since, by pre-training the emulator, it can be applied directly at run-time without retraining on each specific sample. There may also be possibilities for producing ensemble callers by using outputs from multiple existing CNA callers for training.

Our approach could be further improved by explicitly modelling patterns of sequencing variation, due to mappability, amplification or platform sequencing issues. It could also be fine-tuned on real genomes, where profiles have been curated and validated using external data sources (e.g flow cytometry for ploidy estimation, or single cell whole genome sequencing). Further, our modelling approach could likely be extended to integrate sub-clonal calling and holistically account for clone-specific mutations and clone-specific copy numbers. It could likely also be adapted to include multiple longitudinal samples from the same patient, or for low-quality data or different data modalities (e.g SNP array).

A challenge we have highlighted is the issue of identifiability when multiple copy number profiles could explain the same measured sequencing data and the correct copy number profile might be non-identifiable as a result. Multiple tumour sampling in space and/or time, such as used in TracerX (Al Bakir et al., 2023), can alleviate the problem by increasing the probability of unique solutions but may also introduce issues since different evolutionary trajectories could also produce similar patterns of CNAs. In `araCNA` we have addressed this by biasing the training simulations toward lower complexity (ploidy) explanations as is often done with standard CNA callers. We have found that without such bias training data in which there exists a one-tomany relationship between input and output causes `araCNA` to output the average CNA over compatible solutions. Further work is required to enable multiple distinct CNA profiles to be produced.

While deep learning approaches in genomics have previously been limited by a lack of labeled ground truth data and sequence length (Consens et al., 2023), our model `araCNA` demonstrates how simulation-based learning in tandem with emerging deep-learning architectures can address both these issues. We show how `araCNA` performs on par with traditional genomic models despite requiring only a tumour sample and not a matched normal sample, fewer markers of overfitting, and performing inference in only a few minutes. On top of this, `araCNA` has the ability to learn and adapt using fine-tuning or transfer learning.

## Acknowledgements

CY is supported by an UKRI Turing AI Acceleration Fellowship (Ref: EP/V023233/1) and EPSRC grant (Ref: EP/Y018192/1). EV is supported by the Oxford EPSRC Centre for Doctoral Training in Health Data Science (Ref: EP/S02428X/1).

## Data Availability

The latest source codes and tutorials are available on Github (urlhttps://github.com/e-vissch/araCNA). A permanent archive of the software is also available at Figshare (10.6084/m9.figshare.29063492).

