## Supplementary Information for "araCNA: Somatic copy number profiling using long-range sequence models"

### Hyperparameters

Table 1 describes the hyperparameters used for the model architectures of `araCNA` and the simulation-based training.

| Parameter | Description | Value |
|---|---|---|
| **Hyena Backbone Parameters:** unspecified hyperparameters are the same as the defaults provided in Nguyen et al. (2023) | | |
| `d_model` | Projection dimension of data after encoder | 32 |
| `n_layer` | Number of Hyena blocks | 2 |
| `l_max` | Maximum sequence length. Model has capacity for sequences up to this length. Where possible, the model creates arrays of the actual sequence length when less than this to avoid unnecessary memory usage. Some positional encodings of the model cannot be resized and are always set to this value. | 1000000 |
| `causal` | Whether to use bidirectional or causal Hyena blocks. | False |
| `emb_dim` | Dimension of input to inner hyena filter MLP | 5 |
| `w` | Frequency of periodic activations of hyena filter | 10 |
| **Mamba Backbone Parameters:** unspecified hyperparameters are defaults of Mamba package at commit `7fb78a5`, paralleling their `create_blocks` function and `Mamba2` class. | | |
| `d_model` | Projection dimension of data after encoder and backbone. | 32 |
| `n_layer` | Number of bidirectional Mamba2 blocks | 2 |
| `expand` | Dimension of A, B, C parameters in SSM, and the factor by which input dimension expands, before projecting back to model dim. | 4 |
| `headdim` | Dimension of broadcast projection of the input data, before passing through main Mamba2 block. Can be thought of as implementing `headdim` separate SSMs. | 16 |
| `d_intermediate` | Dimension of hidden layer of MLP used as last step in Mamba2 block. If 0, no MLP used. | 0 |
| **Encoder/Embedding Parameters:** | | |
| `input_dim` | Dimension of input data (read depth and BAF) | 2 |
| `embed_dim` | Dimension of embedding, input dimension into backbone | 32 |
| `token_dim` | Number of possible tokens. In `araCNA` only 2 tokens are used- global/not-global, but setting it to 24 allows it to be repurposed in another task with more tokens, e.g chromosome token. | 24 |

**Decoder Parameters:**

| | | |
|---|---|---|
| decoder_dim | Output data dimension of backbone, same as backbone input dimension. | 32 |
| max_tot_cn | Maximum total copy number that can be modelled, total copy numbers higher than this are mapped to a surplus category. | 10 |

**Learning Parameters:**

| | | |
|---|---|---|
| loss_weights | Array corresponding to $\lambda_r, \lambda_p$ for the global reconstruction parameters. | [1, 1] |
| avg_rd_trim_ratio | The proportion of upper/lower quantiles excluded from the average measured read depth in a robust mean calculation. | 0.05 |

**Simulation Parameters:** these correspond to the final simulation parameters, as the simulation difficulty is iteratively increased.

| | | |
|---|---|---|
| read_depth_range | $r_1, r_2$ | [5, 70] |
| purity_range | $\rho_1, \rho_2$ | [0.5, 1] |
| read_depth_scale_range | $\sigma_{r,1}, \sigma_{r,2}$ | [0.01, 0.2] |
| baf_scale_range | $\sigma_{b,1}, \sigma_{b,2}$ | [0.02, 0.1] |
| max_total | The maximum total sampled parental copy number, the sampled minor copy number is less than or equal to the sampled major copy number | 8 |
| max_h_segs | $N_{h,\max}$ | 100 |
| h_l_range | $l_{h,\min}, l_{h,\max}$ | [5, 300] |
| l_min | The minimum segment length, $L_{\min}$. | 100 |
| N | The maximum number of sampled segments, $N$ | max(50, L/1000) |

**Table 1:** Table of Parameters. Some optional parameters in the code base have not been mentioned as they do not affect the model, and are included for possible later development.

### Caller implementations

The following provides more detailed information about the standard CNA callers used in this study for comparison.

#### ASCAT

ASCAT uses a piecewise constant fitting algorithm to segment the data based on step changes present in the read depth (it uses the log ratio between normal and tumour), and BAF (Van Loo et al., 2010). It then uses a grid search approach using values for the purity, $\rho$ and ploidy $\psi$, to find integer copy numbers which best explain the measured data.

ASCAT is implemented according to the to the GitHub, following closely the example given by path `ExampleData/README.md/#ExtractinglogRandBAFfromHTSdataandrunningASCAT`. It requires installation using devtools in R. Please see the araCNA codebase for the exact snakemake workflow used.

ASCAT does not limit the maximum copy number segments are assigned in high read-depth areas, reaching total copy numbers of more than 100 in the TCGA dataset **Figure 6A**. **Supplementary Figure SI-1** investigates the regions captured by these high CN segments.

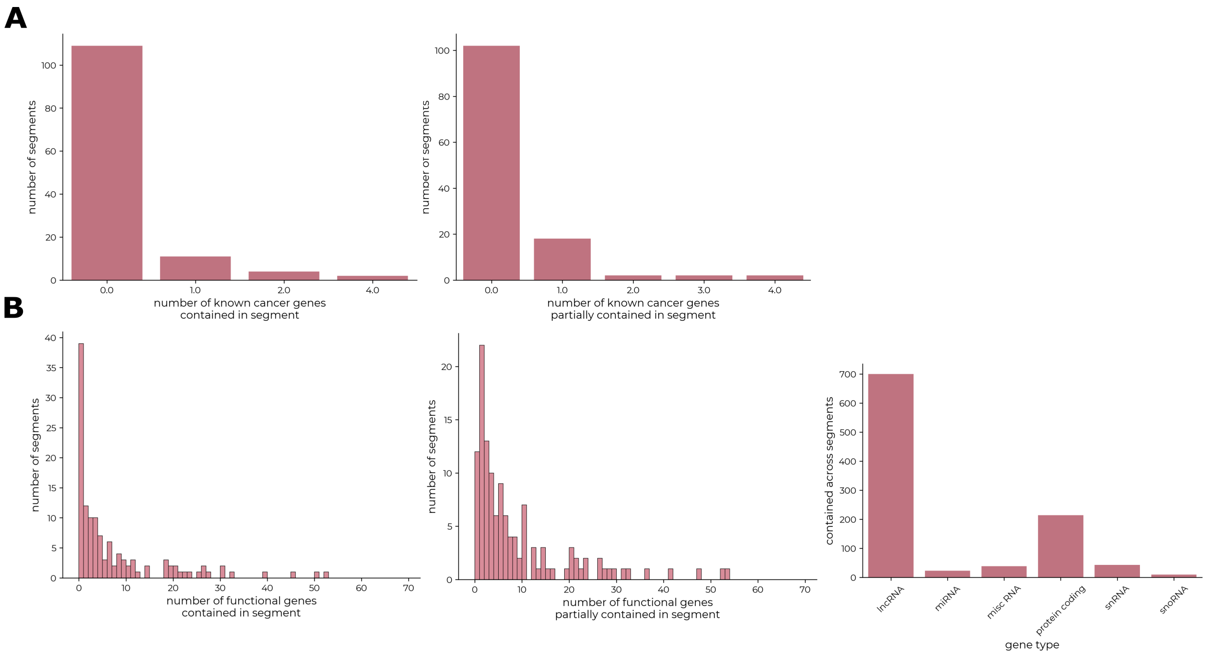


**Figure SI-1: Gene coverage of ASCAT focal aberrations**. The distribution of (A) cancer and (B) all functional genes wholly or partially overlapped by ASCAT focal aberrations (total copy number $\geq 20$). Most ASCAT focal aberrations contain few or no known genes.

#### Battenberg

Battenberg takes a similar approach to ASCAT, however, may also assign a fraction of each segment to a subclone with a different set of copy numbers (Nik-Zainal et al., 2012). This gives an extra set of parameters for each loci/segment; $\tau_i$, the fraction of the sample at that segment attributed to clone 1, $A^1_{M,i}, A^1_{m,i}$, as well as $(1-\tau_i)$, $A^2_{M,i}, A^2_{m,i}$ for clone B (Nik-Zainal et al., 2012). They also add an additional constraint that $A^2_{M,i} + A^2_{m,i} = A^1_{M,i} + A^1_{m,i} \pm 1$, although multiple copies of a region might occur after the emergence of a subclone. If subclones exist in the data, we would expect the distribution $\tau_i$ over segments to be centered around a few distinct modes, where each mode has a different set of copy numbers over regions.

Figure SI-2 indicates that in many samples, the Battenberg multiclone solution is likely overfitting to the input data, as there is a somewhat uniform distribution over the fraction of clone 1, when clones are detected. If clones were to exist in the data, we would expect to see a concentration of fraction values around discrete modes, corresponding to the fraction of cells in a subclone where there exists a diverging copy number pattern across the genome. Battenberg may correctly identify clonal population in case 32 and case 40 where it could be argued there are modes around clonal fractions of 0.5. However, for most other samples, it appears that the Battenberg algorithm is likely overfitting to noisy regions by introducing more modelling parameters than is necessary.

Battenberg is implemented according to the GitHub, following closely example given by path `inst/example/battenberg_wgs.R`. It requires the Bioconductor R package 'battenberg'. Please see the araCNA codebase for the exact snakemake workflow used.

#### HMMCopy

HMMCopy uses only the read depth ratio with normal reads of binned genomic regions, correcting for GC content, and mappability. It uses an HMM to segment the genome into regions of constant copy numbers and call these total copy numbers (Ha et al., 2012).

HMM Copy is implemented according to the manual, which is available with its installation through Bioconductor. It also requires using HMM Copy Utils GitHub for preprocessing the BAM files. The main script follows the Ontario Institute for Cancer Research GitHub script `run_HMMcopy.R`. It should be noted that the HMM parameters sometimes need individual tuning post-inference to obtain fewer total segments and correct the copy number calls (Ha et al., 2012). We adopt the same parameter tuning as the script above.

HMMCopy results could perhaps be improved with further parameter tuning on individual datasets, however, we believe that results obtained without manual tuning capture the reduced utility of this tool compared to other methods that do not require such tuning.

Please see the araCNA codebase for the exact snakemake workflow used, including some preprocessing required as described in the HMM Copy Utils GitHub at `README.md#Example#Onbinwidths`.

#### CNVkit

CNVkit is primarily designed for hybrid capture to infer copy number states, even at regions with very low coverage, however also works for WGS data (Talevich et al., 2016). It uses circular binary segmentation (Venkatraman and Olshen, 2007) to segment the genome into areas of constant copy number, then assigns total copy numbers using thresholds on the log read ratio between normal and tumour. After, it uses BAF to estimate the proportion of each total copy number that can be assigned to each allele (Talevich, 2024). Hence, it only uses the BAF to infer allelic copy numbers after initial total copy number calling.

CNVkit is implemented according to the documentation.

We used the docker image installation, and the 'batch' function pipeline, specifying the method as whole genome sequencing. CNVkit can also do allele-specific calling if provided with VCFs, so we used VarScan to obtain VCFs of the tumour files at SNP loci, followed by calling with CNV kit. CNVkit does not directly account for purity/ploidy but can take both as an input-mainly affecting thresholds for calling copy numbers (Talevich, 2024). Hence, we further implemented CNVkit with ASCAT purity/ploidy as a final comparison point, as this performed better **Figure SI-3**, we have used this, denoted CNVkit*, as the default in the main text.

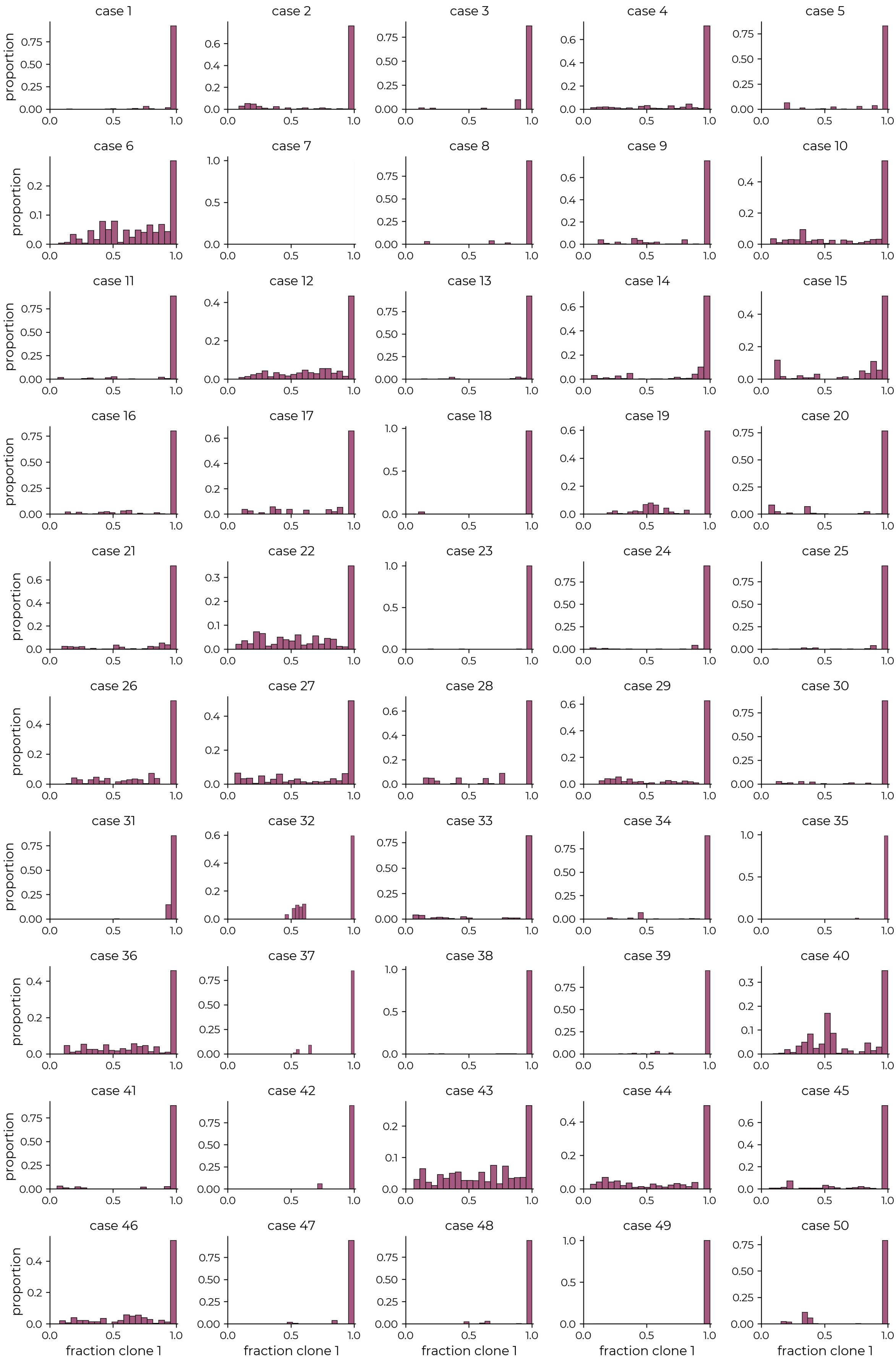


**Figure SI-2:** Distribution of fraction of sample assigned to clone 1 across loci

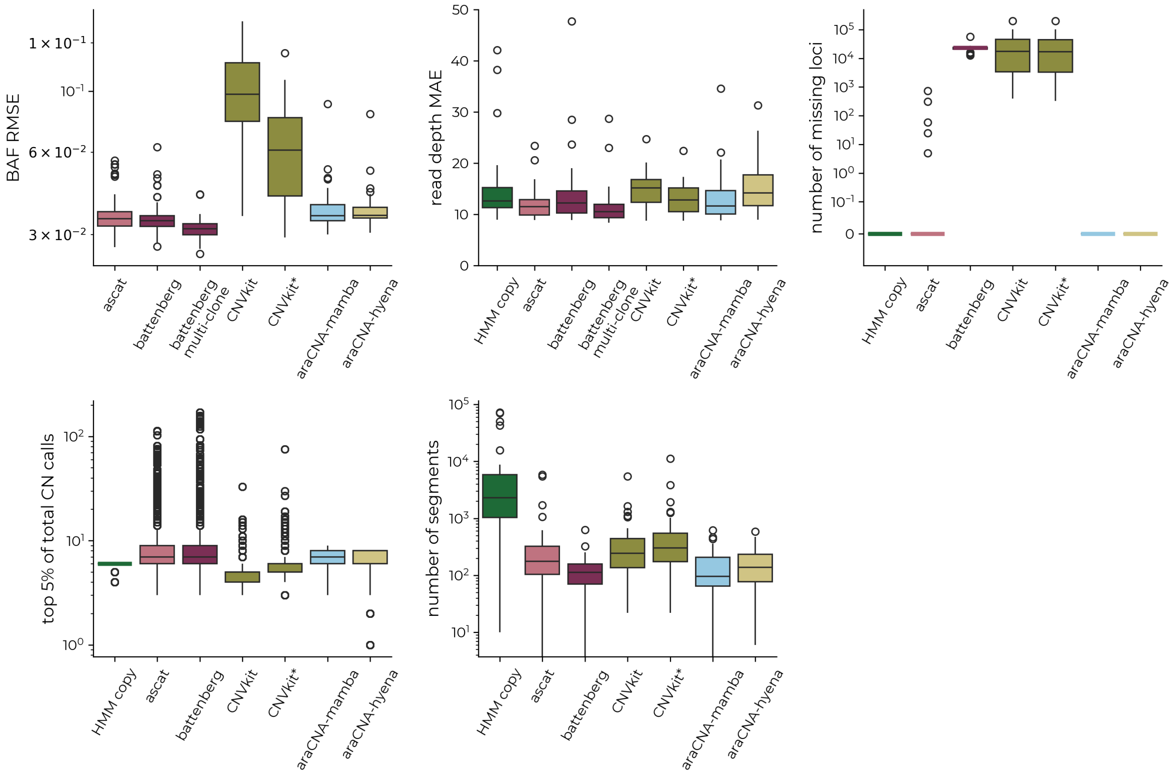


**Figure SI-3:** Summary metrics on 50 TCGA samples demonstrating that CNVkit performs better when provided with the approximate purity- taken from the ASCAT estimates.

### Additional Simulation Details

To summarise the simulation process mathematically, sets of observation data are simulated based on the sampled parental copy number profiles $(A_M, A_m)$ according to the following scheme:

$$\begin{aligned}
\rho &\sim \mathcal{U}[\rho_1, \rho_2], \\
r_d &\sim \mathcal{U}[r_1, r_2], \\
s_{M,i}, s_{m,i} &\sim \text{Bin}(n = 1, p = 0.5), \\
R_i &= r_d C^s_{T,i} + r_d \sigma_r \epsilon_{i,1}, \quad \sigma_r \sim \mathcal{U}[\sigma_{r,1}, \sigma_{r,2}], \quad \epsilon_{i,1} \sim \mathcal{T}_1,
\end{aligned}$$

where $(\rho_1, \rho_2)$, $(r_1, r_2)$, $(\sigma_{r,1}, \sigma_{r,2})$ are hyperparameters and$\mathcal{T}_1$ is Student's t-distribution with 1 degree of freedom.

The BAF sampling is slightly more involved to reflect real data observations, and can be described by the following scheme:

$$\begin{aligned}
n_{h_s} &\sim \mathcal{U}\{1, N_{h,\max}\}, \\
l_{h,j} &\sim \mathcal{U}\{l_{h,\min}, l_{h,\max}\}, \quad S_{h,j} \sim \mathcal{U}\{0, L - l_{h,j}\}, \quad j = 1 \ldots n_{h_s}, \\
N_{r,i} &= \text{Pois}(R_i), \\
N_{B,i} &= \text{Bin}(N_{r,i}, p = \frac{C^s_{B,i}}{C^s_{T,i}}), \\
\nu_i &= \begin{cases} 1, & \text{if } s_{M,i} \neq s_{m,i} \text{ and } i \notin \cup_j \{S_{h,j}, S_{h,j} + l_{h_j}\}, \\ 0.2, & \text{otherwise}, \end{cases} \\
\sigma_b &\sim \mathcal{U}[\sigma_{b,1}, \sigma_{b,2}], \quad \epsilon_{i,2} \sim \mathcal{T}_{150 * \sigma_b}, \\
B'_i &= \begin{cases} \frac{N_{r,i}}{N_{B,i}} + \nu_i \sigma_b \epsilon_{i,2}, & \text{if } 0 \leq \frac{N_{r,i}}{N_{B,i}} + \nu_i \sigma_b \epsilon_2 \leq 1, \\ \frac{N_{r,i}}{N_{B,i}} - \nu_i \sigma_b \epsilon_{i,2}, & \text{otherwise}, \end{cases} \\
B_i &= \max(\min(1, B'_i), 0),
\end{aligned}$$

where $N_{h,\max}$ is the maximum number of homozygous segments, $n_{h_s}$, to sample. $l_{h,\min}$ and $l_{h,\max}$ are the minimum and maximum lengths, $l_{h,j}$ of the homozygous segments, and are hyperparameters, as is $(\sigma_{b,1}, \sigma_{b,2})$. We introduce $N_r$ and $N_B$ as the number of normal and B allele reads, to ensure the frequency is centered around rational values, as would be in real data. $S_{h,j}$ denotes the start positions of the homozygous segments, while $\nu_i$ is the scaling factor that reduces noise at homozygous loci. The parameter $\epsilon_2$ is sampled from a Student's t-distribution with degrees of freedom, df, that scale with the BAF scale parameter- so that smaller noise samples will have a lower df value to ensure that some extreme values are still sampled.
